# EZH2 influences mdDA neuronal differentiation, maintenance and survival

**DOI:** 10.1101/442236

**Authors:** Iris Wever, Lars von Oerthel, Cindy M.R.J. Wagemans, Marten P. Smidt

## Abstract

Over the last decade several components have been identified to be differentially expressed in subsets of mesodiencephalic dopaminergic (mdDA) neurons. These differences in molecular profile have been implied to be involved in the selective degeneration of the SNc neurons in Parkinson’s disease. The emergence and maintenance of individual subsets is dependent on different transcriptional programs already present during development. In addition to the influence of transcription factors, recent studies have led to the hypothesis that modifications of histones might also influence the developmental program of neurons. In this study we focus on the histone methyltransferase EZH2 and its role in the development and maintenance of mdDA neurons. We generated two different conditional knock out (cKO) mice; an *En1Cre* driven cKO, for deletion of *Ezh2* in mdDA progenitors and a *Pitx3Cre* driven cKO, to study the effect of post-mitotic deletion of *Ezh2* on mdDA neurons maturation and neuronal survival. During development *Ezh2* was found to be important for the generation of the proper amount of TH+ neurons. The loss of neurons primarily affected a rostrolateral population, which is also reflected in the analysis of the subset marks, *Ahd2* and *Cck.* In contrast to early genetic ablation, post-mitotic deletion of *Ezh2* did not lead to major developmental defects at E14.5. However, in 6 months old animals *Cck* was found ectopically in the rostral domain of mdDA neurons and *Ahd2* expression was reduced in more mediocaudal positioned cells. In addition, *Pitx3Cre* driven deletion of *Ezh2* led to a progressive loss of TH+ cells in the VTA and these animals display reduced climbing behavior. Together, our data demonstrates that *Ezh2* is important for the generation of mdDA neurons during development and that during adult stages *Ezh2* is important for the preservation of proper neuronal subset identity and survival.

## Introduction

The Substantia Nigra pars compacta (SNc) and the ventral tegmental area (VTA) are two neuronal sub-populations of the mesodiencephalic dopaminergic (mdDA) system, that can already be distinguished during development (Smits et al., 2006; Veenvliet et al., 2013). The projections of the SNc to the dorsolateral Striatum form the nigral-striatal pathway involved in the control of voluntary movement and body posture, while the VTA is involved in the control of emotion-related behavior by innervating the nucleus accumbens, the amygdala and the prefrontal cortex (Prakash and Wurst, 2006; Veenvliet and Smidt, 2014). Even though both groups of neurons use dopamine (DA) as their neurotransmitters, they are molecular distinct and depend on unique transcriptional programs for their development and survival (Veenvliet and Smidt, 2014). Both subsets arise from the same progenitor pool under the influence of differentially expressed transcription factors like *Pitx3* and *En1* (Panman et al., 2014; Veenvliet et al., 2013). Birth-dating experiments showed that neurons of the SNc are born first (Bayer et al., 1995; Bye et al., 2012) and the induction of the SNc phenotype has been shown to be dependent on a complex interplay of transcription factors to antagonize the VTA phenotype. After the establishment of the SNc phenotype it is hypothesized that the remaining DA neurons acquire a VTA phenotype by default (Panman et al., 2014; Veenvliet et al., 2013).

Recent studies have shown that developmental transitions are influenced by Enhancer of Zeste homolog 2 (EZH2) and Polycomb Repressive Complex (PRC) 2 activity (Hirabayashi et al., 2009; Pereira et al., 2010). EZH2 functions as the methyltransferase of the PRC2 complex, which catalyzes the mono- di and tri-methylation of Histone 3 lysine 27 (H3K27) (Cao and Zhang, 2004a; Cao et al., 2002). Methylation of H3K27 negatively influences gene expression by promoting chromatin compaction and shows a highly dynamic profile during development (Cao et al., 2002; Ezhkova et al., 2009; Francis et al., 2004; Margueron et al., 2008; Mohn et al., 2008). Conditional removal of *Ezh2* from cortical progenitors led to a global loss of H3K27 tri-methylation (H3K27me3) and shifted the balance between self-renewal and differentiation, in favor of differentiation (Pereira et al., 2010). A similar role for *Ezh2* was found in neuronal progenitors (NPs) of the dorsal midbrain. *Wnt1Cre* driven deletion of *Ezh2* led to reduced number of NPs in the dorsal midbrain, due to elevated cell cycle exit and differentiation (Zemke et al., 2015). In addition to a role in neuronal development, *Ezh2* has also been linked to several neurodegenerative disorders (Li et al., 2013; von Schimmelmann et al., 2016; Södersten et al., 2014). L-Dopa was found to be capable of negatively influencing the binding of PRC2 to target genes in the Striatum of hemiparkinsonian mice, leading to a de-repression of PRC2 target genes and levodopa-induced dyskinesia (Södersten et al., 2014). In addition, post-mitotic deletion of *Ezh2* in medium spiny neurons and cerebellar purkinje cells, in combination with an *Ezh1* null mutant, led to the derepression of PRC2 targets and a progressive and fatal degeneration of *Ezh2* deficient neurons (von Schimmelmann et al., 2016).

The hallmark of Parkinson’s disease is the specific degeneration of neurons in the SNc, while neurons of the VTA remain largely unaffected (Braak et al., 2003). The specific vulnerability of the SNc neurons is in part thought to be caused by the molecular programming specifics of these neurons. Major progress has been made in unraveling the transcriptional programs involved in the specification and survival of the different subsets of the mdDA system, however little is know about the influence of epigenetics on these processes. In this study we aimed to gain further insight into the role of *Ezh2* in the formation and maintenance of mdDA neurons. To accomplish this we generated two different conditional knock out (cKO) mice; an *En1Cre* driven cKO (Kimmel et al., 2000) for deletion of *Ezh2* in mdDA progenitors and a *Pitx3Cre* driven (Smidt et al., 2012) cKO to study the effects of post-mitotic deletion of *Ezh2* on mdDA neuronal maturation and - survival. Importantly, deletion of *Ezh2* from mdDA progenitors led to a general loss of H3K27me3 in the Cre-recombinant area, while H3K27me3 was still present in cells where *Ezh2* was removed post-mitotically. Analysis of the amount of TH+ cells in developing *En1Cre/+;Ezh2* L/L embryos showed that at E12.5 normal numbers of TH+ cells are generated, however by E14.5 significantly fewer TH+ neurons are detected. The loss of neurons primarily affects the rostrolateral population, which is confirmed through analysis of the subset marks, *Ahd2* and *Cck*. Expression of the rostrolateral mark, *Ahd2*, is significantly reduced in the *En1Cre/Ezh2* cKO, while the expression level of the caudomedial mark, *Cck*, is not affected by the loss of *Ezh2.* In contrast to early genetic ablation, post-mitotic deletion of *Ezh2* did not lead to major alterations in the expression of DA marks at E14.5. However, in 6 months old animals *Cck* was found ectopically in the rostral domain of mdDA neurons and *Ahd2* expression was reduced in more mediocaudal positioned cells. Further analysis of *Pitx3Cre/+; Ezh2* animals demonstrated a progressive loss of TH+ cells in the VTA and reduced climbing behavior in *Pitx3Cre/+; Ezh2* L/L animals. Together, our data demonstrate that *Ezh2* is important for the formation of the population of mdDA neurons during development and that during adult stages *Ezh2* is important for the maintenance of the proper neuronal identity. In addition, our study confirms the initial suggestions that proper *Ezh2* functioning is important for cellular survival, since in our mouse models mdDA neuronal survival is affected and leads to substantial losses.

## Material and Methods

### Ethics Statement

All animal studies are performed in accordance with local animal welfare regulations, as this project has been approved by the animal experimental committee (Dier ethische commissie, Universiteit van Amsterdam; DEC-UvA), and international guidelines.

### Animals

All lines were maintained on a C57BL/6J background (Charles River). *Ezh2*-floxed animals were generated by S.H. Orkin and a kind gift from F. Zilbermann (Friedrich Miescher Institute, Switserland) and have been previously described (Shen et al., 2008). The *En1Cre* line has been generated by A.L Joyner and was a kind gift from S. Blaess (Rheinische Friedrich-Wilhelms-Universität, Gemany) (Kimmel et al., 2000). *En1Cre/+* animals were crossed with *En1Cre-ERT +/+; R R26RYFP/R26RYFP* to obtain *En1Cre/+; R26RYFP/R26RYFP* (Kimmel et al., 2000). The *Pitx3Cre* line has been previously generated in our lab (Smidt et al., 2012). *Ezh2* L/L animals were crossed with *En1Cre/+* or *Pitx3Cre/Cre* animals to obtain *En1Cre/+; Ezh2* L/+ or *Pitx3Cre/Cre; Ezh2* L/+ animals. For the generation of embryos we crosses *En1Cre/+; Ezh2 L/+* or *Pitx3Cre/Cre; Ezh2* L/+ animals with *Ezh2 L/+* animals. Embryos were isolated at embryonic day (E) 12.5, E14.5, considering the morning of plug formation as E0.5. Pregnant or adult mice were euthanized by CO2 asphyxitian and embryos or brains were collected in 1xPBS and immediately frozen on dry-ice (fresh frozen) or fixed by immersion in 4% paraformaldehyde (PFA) for 4-8hrs at 4°C. After PFA incubation, samples were cryoprotected O/N in 30% sucrose at 4°C. Embryos or brains were frozen on dry-ice and stored at −80°C. Cryosections were slices at 16μm, mounted at Superfrost plus slides, air-dried and stored at −80°C until further use.

### Genotyping

The genotyping for the *Ezh2-flox* allele was executed with 50-100 ng of genomic DNA together with forward primer 5’-ACCATGTGCTGAAACCAACAG -3’ and reverse primer 5’-TGACATGGGCCTCATAGTGAC – 3’ resulting in a 395 bp product for the wildtype allele and a 361 bp product for *floxed* allele.

Genotyping of the *En1Cre* allele was performed with 50-100 ng of genomic DNA together with primer pair En1Cre 5UTR_F3 5’-CTTCGCTGAGGCTTCGCTTT-3’ and En1Cre Cre_R2 5’-AGTTTTTACTGCCAGACCGC-3’ resulting in a product at 240 bp for the *Cre*-allele.

Pitx3-Cre genotyping was done by two different PCR’s using 50-100ng of genomic DNA for both reactions. The mutant allele was detected by using primer pair forward 5’-GCATGATTTCAGGGATGGAC and reverse 5’-ATGCTCCTGTCTGTGTGCAG, resulting in a product of 750 bp for a mutant allele, and no product in wild-type animals. To additionally detect the wildtype allele primers were designed around Pitx3 exon 2 and exon 3’ forward 5’-CAAGGGGCAGGAGCACA and reverse 5’-GTGAGGTTGGTCCACACCG, resulting in a product of 390 bp for the wildtype allele and no product for the mutant allele.

Genotyping for the *R26R-YFP* allele was performed using 3 primers Rosa_mutant primer 5’- AAGACCGCGAAGAGTTTGTC -3’, Rosa_wildtype primer 5’- GGAGCGGGAGAAATGGATATG -3’ and a Rosa_common primer 5’-AAAGTCGCTCTGAGTTGTTAT - 3’ with 50-100 ng of genomic DNA. The PCR reaction gave a product at 320 bp for the mutant *R26R-YFP* allele and a product of 600 bp for the wildtype allele.

### In situ hybridization

*In situ* hybridization with digoxigenin (DIG)-labeled RNA probes was performed as described (Smidt et al., 2004; Smits et al., 2003). DIG-Labeled RNA probes for *Th, Ahd2, Cck* and *Dat* have been respectively described (Grima et al., 1985; Jacobs et al., 2007, 2011; Smits et al., 2003). The Calb1 probe was a 509bp fragment directed against bp 196-704 of the Calbindin D 28K mRNA (NM_009788).

### Immunohistochemistry

#### Fluorescent immunohistochemistry

Cryosections were blocked with 4% HiFCS in 1x THZT [50mM Tris-HCL, pH 7.6; 0.5M NaCl; 0.5% Triton X-100] and incubated with a primary antibody [Rb-TH (Pelfreeze 1:1000), Sh-TH (Millipore AB1542, 1:750), Rb-H3K27me3 (Millipore, 17-622 1:2000), Rb-Pitx3 (1:750, (Smidt et al., 2000)] in 1xTHZT overnight at room temperature. The following day the slides were washed and incubated for 2 hrs at room temperature with secondary Alexafluor antibody (anti-rabbit, anti-sheep (In vitrogen, 1:1000) in 1xTBS. After immunostaining nuclei were staining with DAPI (1:3000) and embedded in Fluorsave (Calbiogen) and analyzed with the use of a fluorescent microscope (Leica). All washes were done in TBS and double stainings were performed sequential, with immunostaining for TH being done first, followed by the staining for H3K27me3. The antibody against H3K27me3 requires antigen retrieval, which was executed as follows; slides were incubates 10 minutes in PFA after which they were submersed in 0.1M citrate buffer pH 6.0 for 3 minutes at 1000Watts followed by 9 minutes at 200Watts. Slides were left to cool down, after which protocol was followed as described above.

### Quantitative analysis

Quantification of TH-expressing neurons 3 month old and 6 month old midbrain was performed in ImageJ as follows. Cells were counted in 10-12 matching coronal sections (3 months old; n=3, 6 months old; n=3). Cells were counted as TH+neurons when TH staining co-localized with nuclear DAPI staining. The separation of the SNc and VTA were made based on anatomical landmarks. Everything rostral of the supramammillary decussation was considered as SNc and distinction between the SNc and the VTA was made based on the tracts medial lemniscus, positioned between the SNc and VTA. Quantification of the TH-expressing cells in embryos was performed in ImageJ with particle analysis. TH (green) and DAPI (blue) images were turned into binary images using default settings, after which the binary DAPI image was used as an overlay on top of the TH image via image calculator ‘and’. This generates a binary image of only the cells that are positive for both TH and DAPI, which were then counted by using the ‘analyze particles’ function of ImageJ. For E12.5 10-14 matching sagittal sections were analysed and for E14.5 15-17 matching sagittal sections were analyzed. The separation between medial and lateral was made based on the retroflex fasciculus. All sections, including the section portraying the retroflex fasciculus were considered lateral. Statistical analysis was performed via a student’s T-test.

### Quantitative PCR (qPCR)

RNA was isolated from dissected E14.5 midbrains of *En1Cre/+; Ezh2 +/+* and *Ezh2* cKO embryos. RNA was isolated with Trizol (ThermoFisher) according to manufacturers protocol. Two midbrains were pooled for the *Ezh2* cKO samples and a single midbrain was used per sample for the wildtype (Wildtype n=4 and *En1Cre/+; Ezh2* L/L n=3). Relative expression levels were determined by using the QuantiTect SYBR green PCR lightCycler kit (Qiagen) according to the manufacturers instructions. For each reaction 10ng (dissected midbrain) of total RNA was used as input. Primers used for *Th, Ahd2, Cck* and were previously published (Jacobs et al., 2011). Primers for *Pitx3* were designed as follows: forward 5’ - CTTCCAGAGGAATCGCTACCCT and reverse 5’-CTGCGAAGCCACCTTTGCACA (product size 164bp).

### Behavioral analysis

#### Climbing test

Climbing behavior was assessed as described before (Protais et al., 1976; Smidt et al., 2004) during the dark (active) phase between 21.00 and 23.00. Animals were assessed twice, once at 3 months and once at 6 months. Animals were placed in the climbing cylinders and acclimatized for 30 minutes. All behavioral observations were done in a separate behavioral room to which the animals were transported 1 hr prior to the experiment.

#### Open field test

The open field consisted of a plastic open rectangular box (54 × 37 × 33 cm) with bedding material on the bottom. Locomotor activity was monitored for 15 minutes using a fully automated observation system (Ethovision, Noldus Information Technology, The Netherlands). The animals were tested twice, once at 3 months and once at 6 months of age. Measurements were performed during the dark (active) phase between 21.00 and 23.00 and in a separate behavioral room to which the animals were transferred 1 hr prior to the experiment.

## Results

### H3K27me3 is lost in the *Cre*-recombinant area of *En1Cre* driven *Ezh2* cKOs

Previous studies have shown that the conditional deletion of *Ezh2* in neuronal progenitors shift the balance between self-renewal and differentiation, in favor of differentiation (Feng et al., 2016; Pereira et al., 2010; Zemke et al., 2015). Conditional deletion of *Ezh2* in cortical progenitors showed that an increased fraction of cortical progenitors leave the cell cycle at an earlier time-point, leading to a substantial thinner cortex (Pereira et al., 2010). Matching results were obtained for the dorsal midbrain, where the loss of *Ezh2* leads to a reduced neuroepithelial thickness, by negatively affecting proliferation and canonical Wnt-signaling (Zemke et al., 2015). To study whether *Ezh2* affects the differentiation of mdDA neurons we crossed Ezh2-floxed mice (Shen et al., 2008) with *En1Cre* animals (Kimmel et al., 2000), deleting *Ezh2* from ventral midbrain progenitors from E10.5 onward (Sunmonu et al., 2009; Zemke et al., 2015). As described above, EZH2 is the methyltransferase of PRC2, which catalyzes the methylation of H3K27 (Cao and Zhang, 2004a; Margueron et al., 2008; Shen et al., 2008) and in previous studies in which *Ezh2* is conditionally removed from neuronal progenitors a widespread loss of H3K27me3 is observed (Hirabayashi et al., 2009; Pereira et al., 2010; Zemke et al., 2015). To determine whether *En1Cre* driven deletion of *Ezh2* affects the presence of H3K27me3 we performed immunohistochemistry experiments for H3K27Me3 at E14.5. The Cre-recombinant area was visualized using the R26R-YFP reporter allele (Srinivas et al., 2001) and in accordance with previous studies, H3K27me3 is lost in the Cre-recombinant area of *En1Cre/+;Ezh2* L/L; R26R-YFP/ R26R-YFP embryos (**Figure 1 (1,** right panel)), while the mark is still present in wildtypes (**Figure 1 (1,** left panel)). Analysis of the periphery of the *En1Cre* expression domain demonstrated a mixture of H3K27me3 positive and negative cells in these *Ezh2* cKO embryos (**Figure 1(2,** right panel), arrowhead), suggesting that the recombination at the border regions is incomplete. Together, our data shows that the early removal of *Ezh2* is sufficient to disturb general PRC2 functioning and ablates the tri-methylation of H3K27 in the *Cre*-recombinant area.

**Figure 1:**
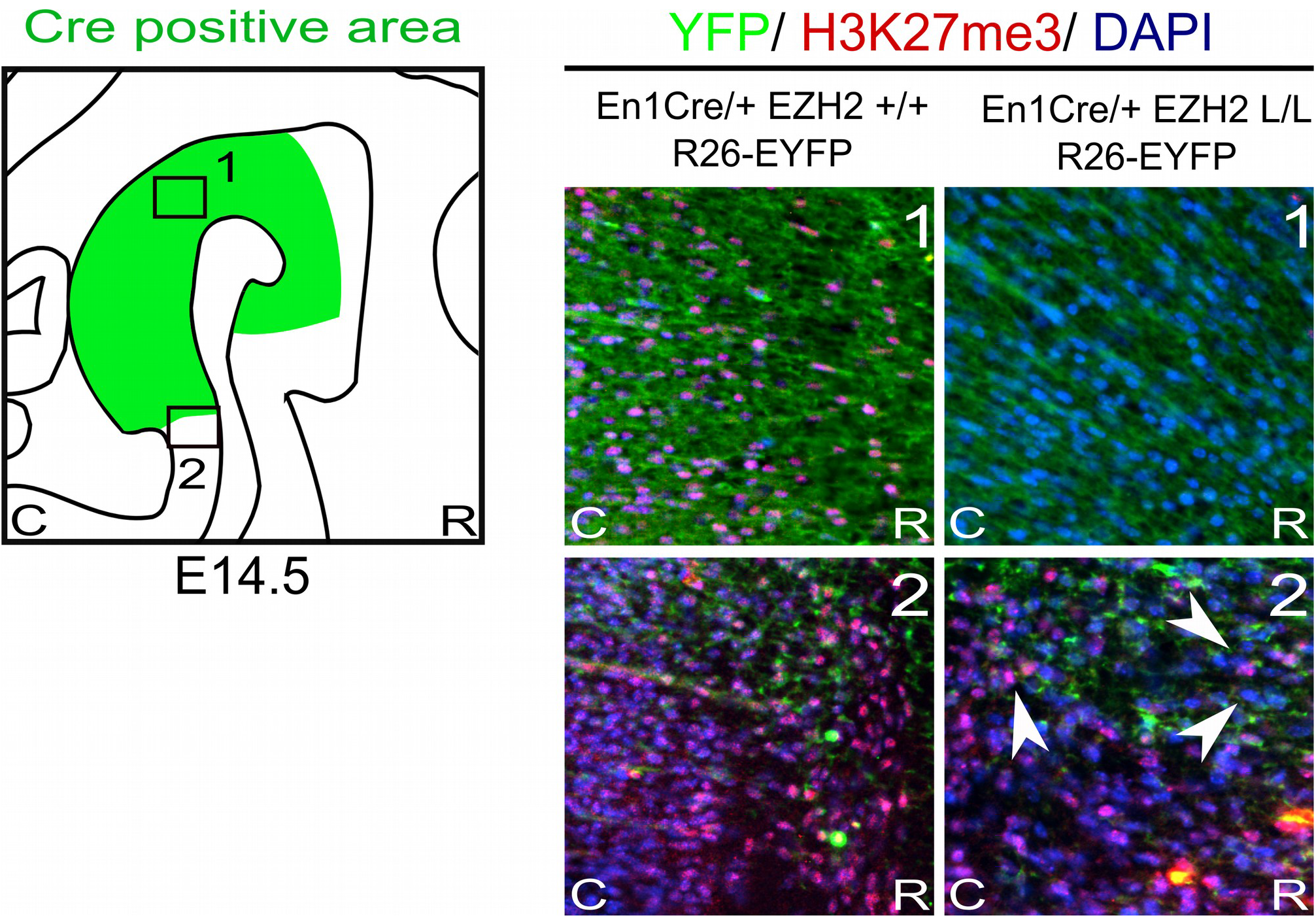
H3K27me3 is lost in the Cre-recombinant area of *En1Cre* driven *Ezh2* cKO embryos. Analysis of the presence of H3K27me3 (red) in the *Cre* recombinant area, marked by YFP (green), in *En1Cre/+; Ezh2* +/+ and *En1Cre/+; Ezh2* L/L E14.5 embryos. In *En1Cre/+; Ezh2* L/L embryos H3K27me3 could not be detected in the midbrain at E14.5, while it was present in wildtype control embryos (1). Outside of the Cre-recombinant area H3K27me3 was present in both the wildtype and *En1Cre/Ezh2* mutant (2). Co-localization with DAPI staining was used to confirm nuclear staining.

### *En1Cre/+; Ezh2* L/L embryos display a disorganized mdDA domain and a reduction in TH positive neurons

MdDA progenitors develop at the ventricular zone of the ventral midbrain (Brodski et al., 2003; Mesman et al., 2014; Ono et al., 2007; Placzek and Briscoe, 2005). Around embryonic day (E) 10.5 the first progenitors exit the cell cycle to give rise to post-mitotic mdDA precursors that will start to express the rate-limiting enzyme of DA synthesis, Tyrosine Hydroxylase (TH) (Bayer et al., 1995). To study the effect of *En1Cre* driven deletion of *Ezh2* on mdDA neurogenesis we quantified the amount of TH+ neurons at E12.5, at the peak of neurogenesis (Bayer et al., 1995; Bye et al., 2012), and E14.5, before the first wave of apoptosis commences (Bayer et al., 1995; Bye et al., 2012; Zhang et al., 2007), by performing immunohistochemistry for TH (**Figure 2**). Spatial analysis of the TH+ domain at E12.5 did not reveal any changes in the DA region between *En1Cre/+; Ezh2 +/+* and *En1Cre/+;Ezh2* L/L embryos. Also quantification of the amount of TH+ cells did not show any significant differences between the *Ezh2* cKO and wildtype animals (**Figure 2A**). However when examining the total number of TH+ neurons at E14.5 a loss of TH+ cells was found in the *En1Cre/ +; Ezh2* L/L midbrain (n=3, p<0.05, two-tailed) (**Figure 2B**).

**Figure 2:**
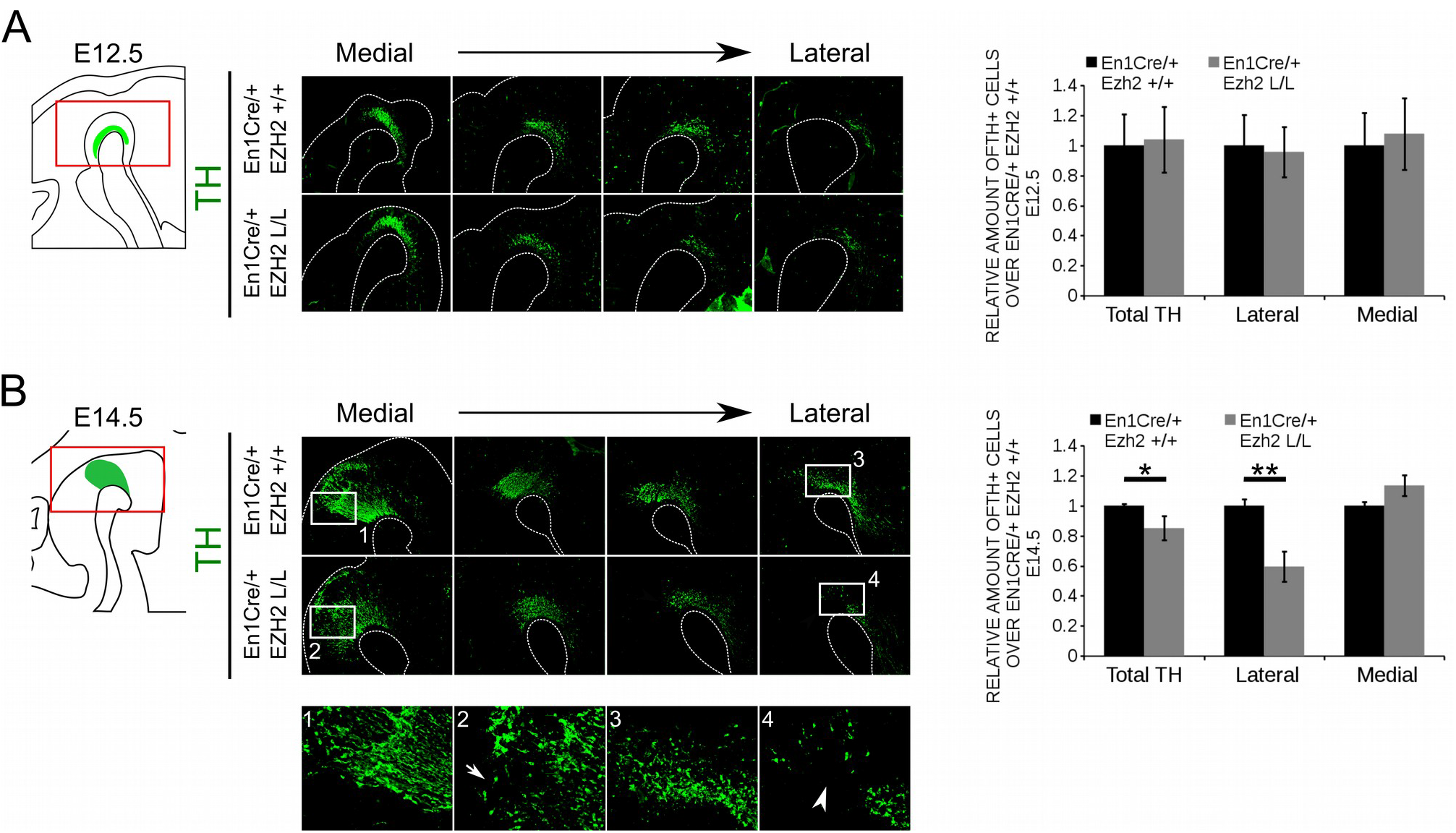
*En1Cre* driven deletion of *Ezh2* affects the amount of TH+ cells at E14.5. Protein expression of TH was evaluated at E12.5 (A) and E14.5 (B). (A) Quantification of the amount of TH positive cells in sagittal midbrain sections at E12.5 demonstrated no significant changes in the number of TH+ cells between the *En1Cre/+;Ezh2* +/+ (n=3,black bar) and *En1Cre/+Ezh2* L/L (n=3, grey bar) embryos. Wildtypes were set at 1. In addition TH+ neurons were normally distributed over the lateral and medial mdDA domain. (B) Quantification of the amount of TH+ in sagittal midbrain sections at E14.5. The total amount of TH+ neurons were reduced with ∼15% in the *Ezh2* cKO embryos (n=3, *P<0.05, two-tailed). Subdivision into the lateral and medial population based on the location of the retroflex fasciculus demonstrates that in the lateral population ∼ 41% of the TH+ cells are lost in *En1Cre/+; Ezh2* L/L embryos (4, white arrowhead) (n=3, **P<0.01, two-tailed), while an upward trend in the amount of TH+ cells in the medial domain is observed (2, arrow) (n=3, P= 0.089, two-tailed). Wildtypes were set at 1.

During neurogenesis post-mitotic mdDA cells migrate via radial and tangential migration to their final location to form different subsets of mdDA neurons (Kawano et al., 1995; Shults et al., 1990; Smits et al., 2013). While neurons destined to become the SNc are predominantly found in the rostrolateral population, the neurons that form the VTA are located caudomedial (Veenvliet et al., 2013). When examining the expression pattern of TH, we observed that mostly the rostrolateral population was affected in *En1Cre/+; Ezh2* L/L animals (**Figure 2B (4)**, arrowhead). Quantification of the amount of cells found rostrolateral and caudomedial showed that ∼ 40% of the rostrolateral neurons were lost (n=3, p < 0.01, two-tailed), while an upward trend in the number of TH+ neurons located caudomedial was found (**Figure 2B (2)**) (n=3, p=0.08, two-tailed), suggesting that, next to a loss, TH+ neurons may be dislocated. In agreement with sagittal sections, the most extensive loss of TH+ neurons in coronal sections was observed in rostral sections of *En1Cre/Ezh2* mutant embryos (**Supplemental 1**, arrowheads). Together, these data show that *Ezh2* is important for the generation of the proper amount of TH+ neurons at the proper positions and that the loss of *Ezh2* mainly affects the rostrolateral population of TH+ cells.

### MdDA subsets are differently affected by *En1Cre* driven deletion of *Ezh2*

As described above, different mdDA subsets can already be distinguished during embryonic development based on their anatomical location (La Manno et al., 2016; Smits et al., 2013; Veenvliet et al., 2013). In addition to their location, each mdDA subset is characterized by a unique molecular code (Di Giovannantonio et al., 2013; Di Salvio et al., 2010; La Manno et al., 2016; Panman et al., 2014; Veenvliet et al., 2013), with *Ahd2* and *Cck* as hallmarks for the rostrolateral and caudomedial populations respectively (Jacobs et al., 2007; Veenvliet et al., 2013). Since *En1Cre* driven deletion of *Ezh2* leads to a major loss of TH+ cells in the rostrolateral population, we aimed to confirm this by analyzing the expression of these subsets marks. The expression of *Th, Cck* and *Ahd2* was analyzed using *in situ* hybridization and the levels were quantified by qPCR (**Figure 3**). In accordance with TH protein data, *Th* expression was affected in rostrolateral sections of E14.5 *En1Cre/Ezh2* mutant midbrains (**Figure 3A** arrowhead). Noteworthy, the overall expression level was not significantly altered (**Figure 3B**). Analysis of two subset marks, *Ahd2* and *Cck*, demonstrated that the overall expression level of the rostrolateral mark *Ahd2* was reduced to ∼27% in *En1Cre/+; Ezh2* L/L animals (n=3) (**Figure 3B**) in comparison to wildtypes (n=4)(p<0.05, two-tailed) and spatial expression analysis showed that signal was lost in both lateral and medial sections of the midbrain of *En1Cre* driven *Ezh2* cKO embryos (**Figure 3A**, arrowheads). In contrast, *Cck* expression was found to be extended rostrally in the medial-ventral-midbrain in *En1Cre/+; Ezh2* L/L embryos (**Figure 3A**, arrows). Again, the overall expression level was comparable to wildtypes (**Figure 3B**).

**Figure 3:**
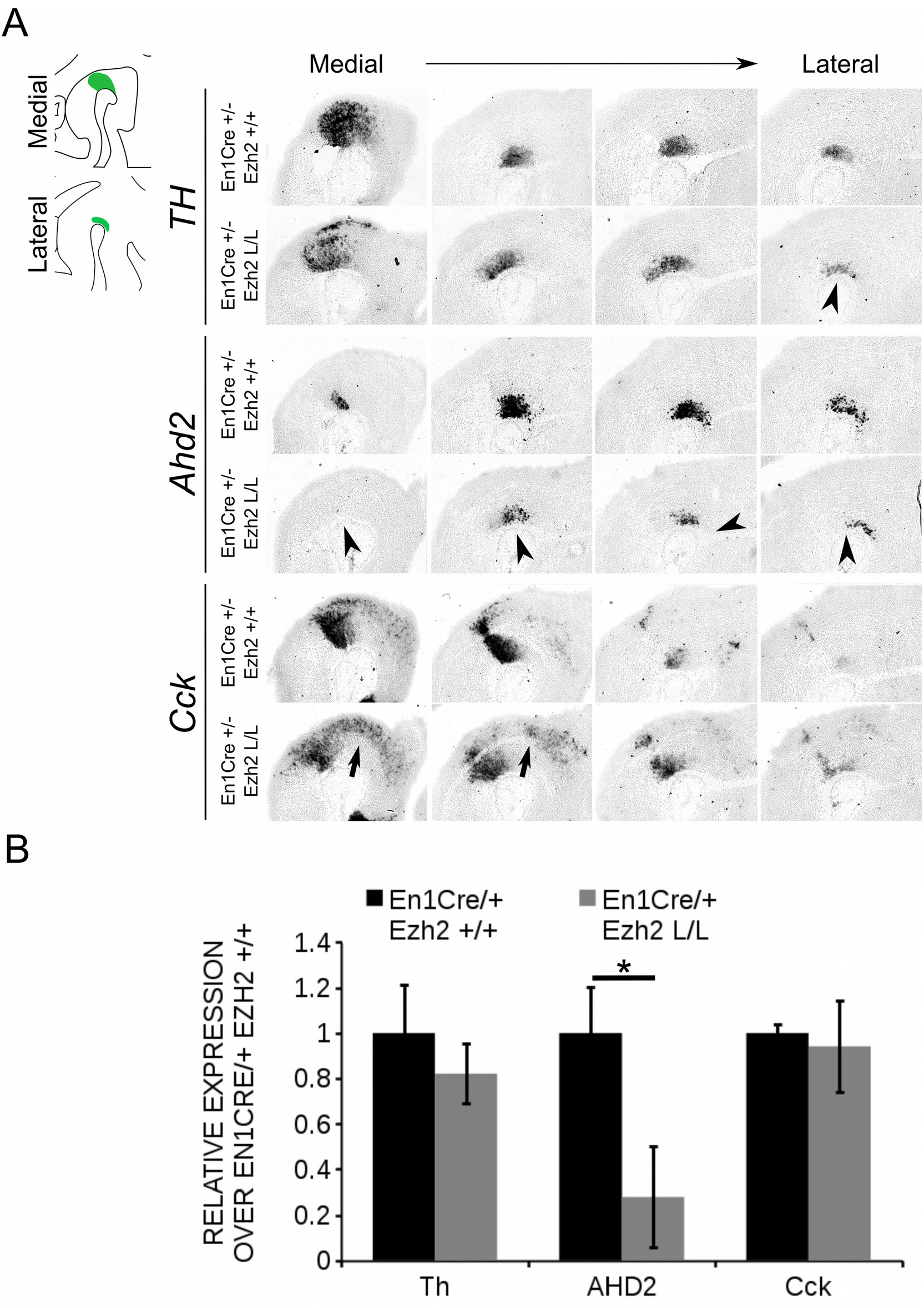
*Ahd2* expression is reduced in *En1Cre/+; Ezh2* L/L embryos at E14.5. (A) *In situ* hybridization for *Th, Cck* and *Ahd2* in sagittal E14.5 midbrain sections. *Ahd2* expression is reduced in both lateral en medial sections (lowest panel, arrowheads), while *Th* expression is decreased only in the lateral sections (upper panel, arrowhead). Expression of *Cck* was found ectopically in the rostral midbrain (arrows). (B) Quantification of the expression levels of *Th, Cck* and *Ahd2* via qPCR showed that *Th* and *Cck* levels were not significantly altered in the *En1Cre/+; Ezh2* L/L (grey bars), but levels of *Ahd2* were reduced to ∼20% of the wildtype level (black bar) (n=3, *P<0.05, two-tailed). Wildtype levels were set at 1.

The phenotype observed in *En1Cre/+; Ezh2* L/L embryos is partially reminiscent of defects observed in *Pitx3* mutants (Jacobs et al., 2007, 2011), in which *Ahd2* expression is lost and expression of *Cck* is up-regulated and expanded into the rostrolateral mdDA population (Jacobs et al., 2007, 2011). To verify whether the programming deficiency observed in *En1Cre/+; Ezh2* L/L animals is not due to the possible lack of *Pitx3* we performed immunohistochemistry for PITX3 at E14.5 and quantified the mRNA levels by means of qPCR (**Figure 4**). PITX3 was detected in lateral and medial midbrain sections of both *En1Cre/+; Ezh2* +/+ and *En1Cre/+; Ezh2* L/L embryos and mimicked the expression pattern of TH (**Figure 4A**). Quantification of the mRNA levels showed that the *Pitx3* levels are not significantly different between *En1Cre/+; Ezh2* L/L and control animals (**Figure 4B**), indicating that the observed phenotype in *En1Cre* driven *Ezh2* cKOs is not due to any *Pitx3* deficiency. In summary, our data show that a loss of *Ezh2* affects the positioning of the *Th* population and differently affects mdDA subsets, leading to the loss of rostrolateral *Ahd2* expression and extension of the *Cck* expression domain.

**Figure 4:**
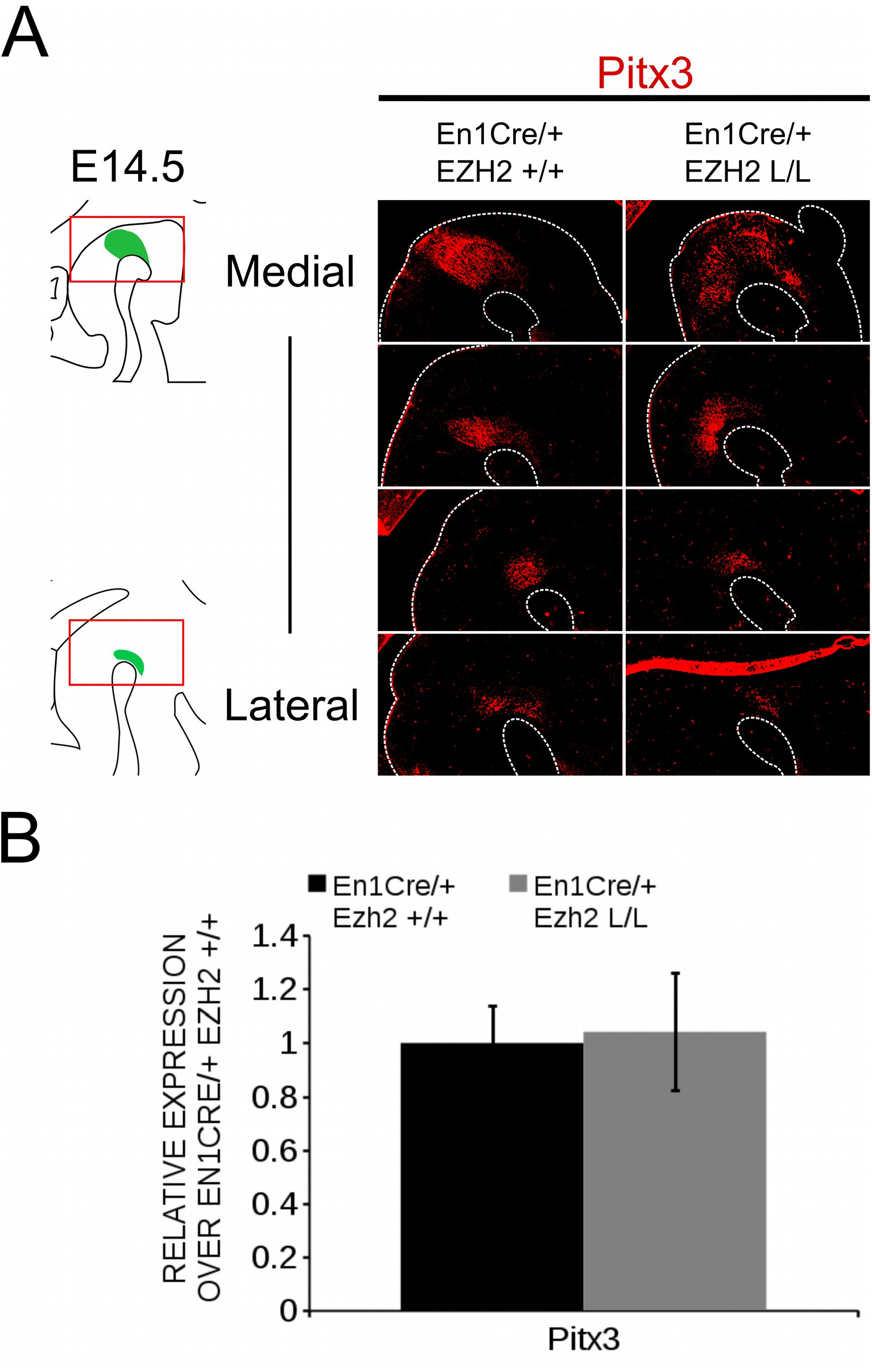
Pitx3 expression is not influenced by *En1Cre* driven deletion of *Ezh2* at E14.5. (A) Analysis of the PITX3 protein expression in E14.5 *En1Cre/+; EZh2* +/+ and *En1Cre/+;Ezh2* L/L midbrains by means of immunohistochemistry. PITX3 expression can be detected in both the medial and lateral sections of the midbrain of Ezh2_cKO embryos, but less PITX3 + cells are present in the lateral sections of *En1Cre/+; Ezh2* L/L embryos in comparison to wildtype littermates (white arrowheads). (B) Quantification of the mRNA levels by means of qPCR demonstrated not significant changes in expression levels between the *En1Cre/+; Ezh2* +/+ (black bar) and the *Ezh2* cKO (grey bar) (n=3). Wildtype levels where set at 1.

### Post-mitotic deletion of *Ezh2* affects the expression of subset specific factors *Cck* and *Ahd2* in 6 month old animals

Next to early developmental influences, we aimed to study the effect of *Ezh2* deletion on mdDA neuronal maturation. However, similar to *Wnt1Cre/+;Ezh2* mutants (Zemke et al., 2015), *En1Cre* driven deletion of *Ezh2* leads to prenatal lethality. We therefore generated a second model, in which *Ezh2* was deleted specifically in post-mitotic mdDA progenitors by crossing the Ezh2-floxed line (Shen et al., 2008) with *Pitx3Cre* animals (Smidt et al., 2012), to study mdDA neuronal maturation. To substantiate the initial phenotype found in *En1Cre/Ezh2* mutants we examined the mdDA markers *Th, Ahd2* and *Cck* in *Pitx3/Ezh2* mutants. At E14.5 no differences in *Th* expression could be observed between *Pitx3Cre/+;Ezh2* +/+ and *Pitx3Cre/+; Ezh2* L/L embryos (**Figure 5**).

**Figure 5:**
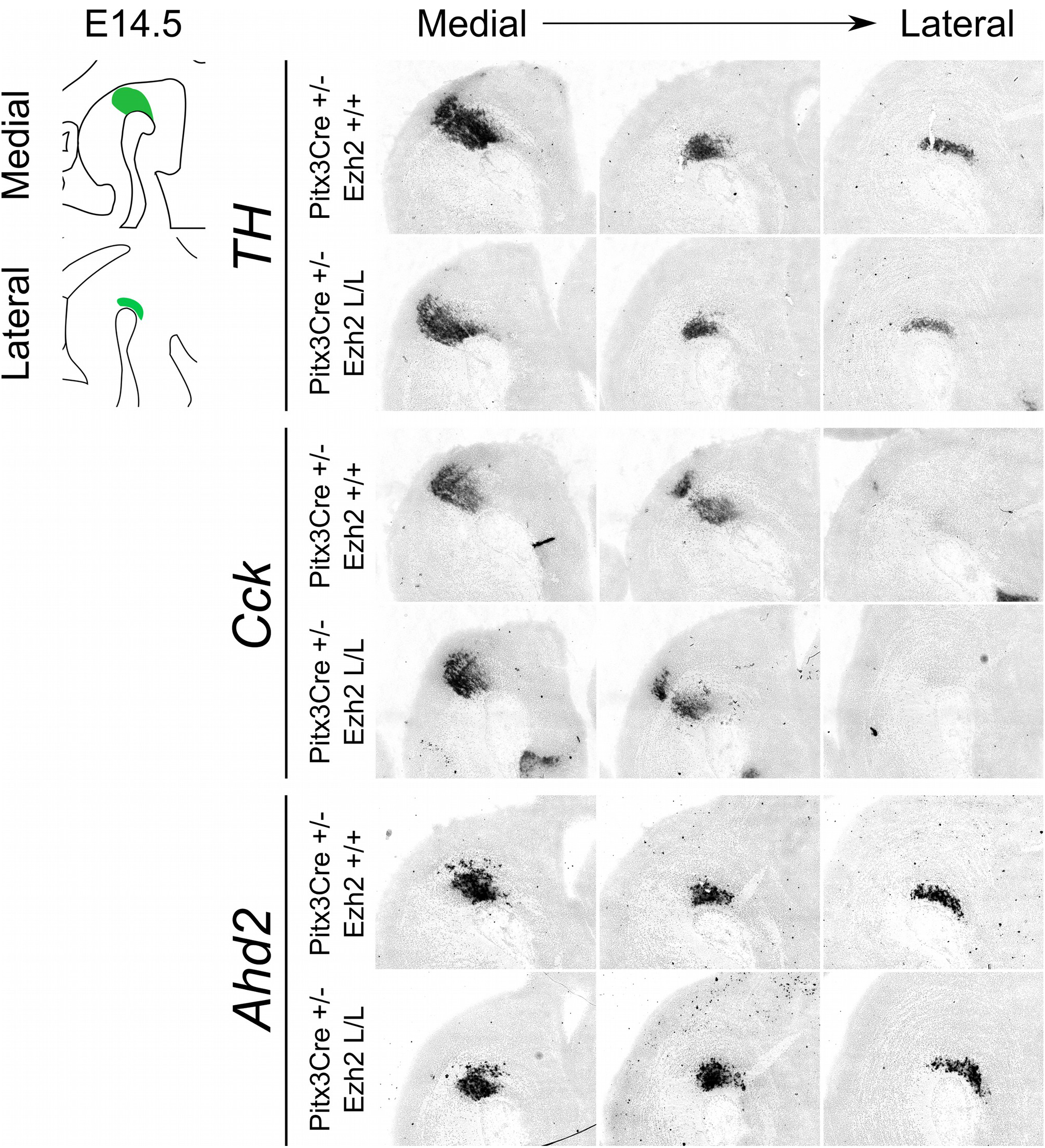
Post-mitotic deletion of *Ezh2* does not change the expression pattern of *Cck* and *Ahd2* at E14.5. Spatial expression of *Th, Cck* and *Ahd2* were analyzed in E14.5 sagittal midbrain sections of *Pitx3Cre/+; Ezh2 +/+* and *Pitx3Cre/+: Ezh2* L/L embryos with *in situ* hybridization. *Th, Cck* and *Ahd2* all show a normal spatial expression in both lateral and medial sections in the *Ezh2* cKO in comparison to the wildtype sections.

Moreover, *Ahd2* and *Cck* marked the default distinct mdDA sub-domains (Veenvliet et al., 2013) in the sagittal midbrain of both wildtype and *Pitx3Cre/+; Ezh2* L/L embryos (**Figure 5**). After the analysis of the embryonic stage, we proceeded with examining the expression of *Th, Cck* and *Ahd2* in the midbrain of 3 months and 6 months old *Pitx3Cre/+; Ezh2* L/L animals (**Figure 6**).

**Figure 6:**
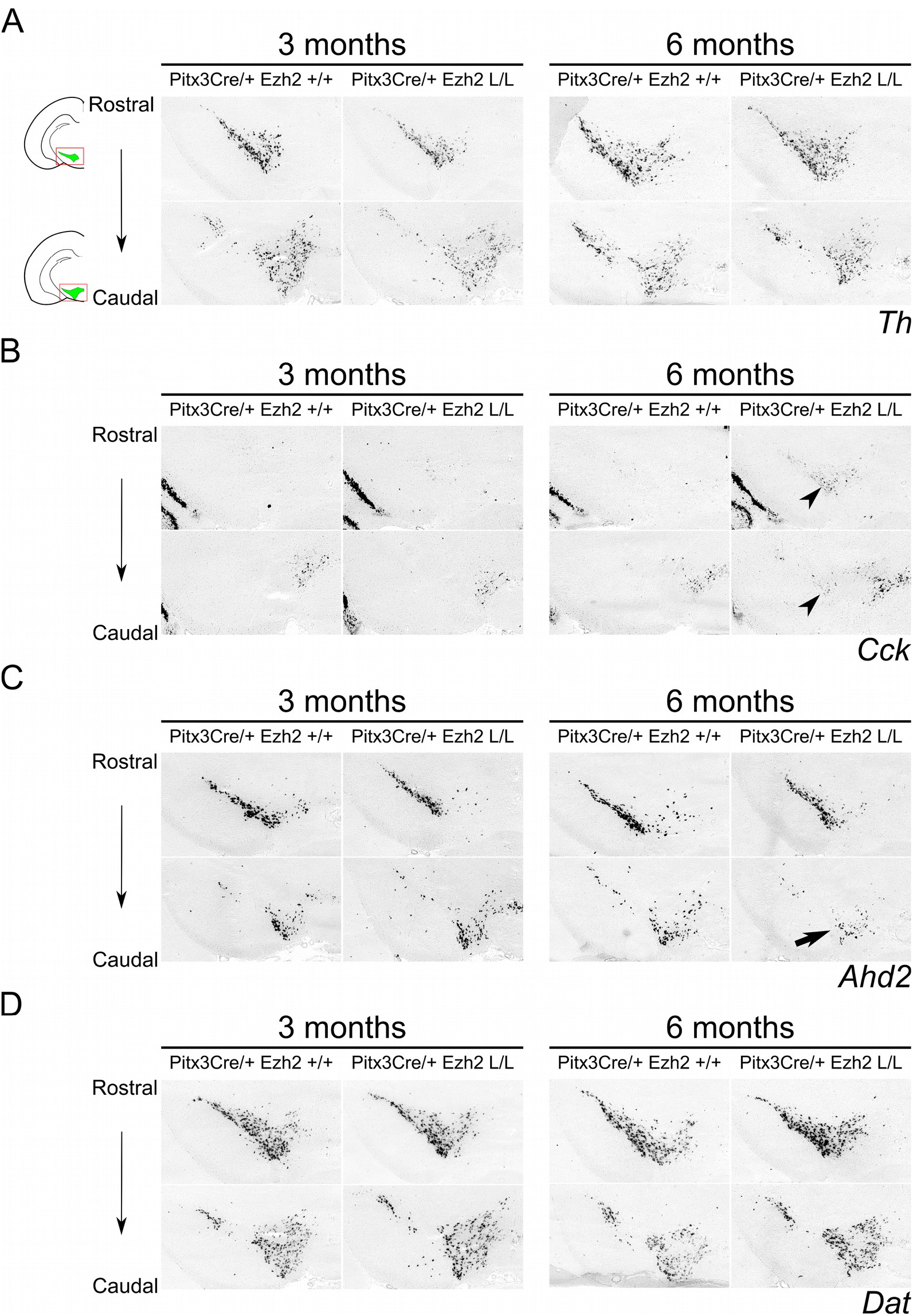
Expression of *Cck* and *Ahd2* is altered in the midbrain of 6 month old *Pitx3Cre/+; Ezh2* L/L animals. *In situ* hybridization of *Th, Dat, Cck*, and *Ahd2* in 3 month old and 6 month old wildtype and *Ezh2* cKO animals. (A) Analysis expression pattern of the general DA mark *Th* does not reveal considerable changes in expression between the *Pitx3Cre/+; Ezh2* +/+ and the *Ezh2* cKO. (B) *Cck* is ectopically expressed in the rostral sections of the midbrain of 6 month old *Pitx3Cre/+; Ezh2* L/L animals (arrowheads). This rostral expansion of *Cck* expression is not observed in 3 month old *Ezh2* cKO animals (left panel). (V) Expression of *Ahd2* is progressively reduced in the caudal midbrain of 6 month old *Pitx3Cre/+; Ezh2* L/L animals in comparison to wildtype and 3 month old *Ezh2* cKO animals (arrow), while expression in the rostral sections is maintained. (D) The expression pattern of *Dat* in *Pitx3Cre/+;Ezh2* L/L adult midbrain is similar to wildtype littermates at both 3 months and 6 months of age.

Examination of the expression of *Th* demonstrated no clear expression pattern differences between *Pitx3Cre/+; Ezh2* +/+ and *Pitx3Cre/+; Ezh2* L/L genotypes (**Figure 6A**). Assessment of the expression pattern of both *Cck* and *Ahd2* revealed that post-mitotic deletion of *Ezh2* influences the expression of both marks in a similar direction as early deletion of *Ezh2* alters *Cck* and *Ahd2* expression (**Figure 2; 6B, C**). In 6 month old *Ezh2* cKO animals ectopic expression of *Cck* was found in the rostrolateral domain of mdDA neurons (**Figure 6B**, arrowheads), while *Ahd2* expression was reduced in caudal areas (**Figure 6C**, arrow). Interestingly, the expression of *Cck* and *Ahd2* was unaffected in 3 month old *Pitx3Cre/+; Ezh2* L/L animals and *Ahd2* was also normally expressed in the rostral domain in 6 month old mutants (**Figure 6B, C**). Next to these mdDA markers, we we also included the expression pattern of the *Dopamine transporter* (*Dat*) in the adult midbrain (**Figure 6D**). During development *Dat* expression partially overlaps with the expression domains of both *Ahd2* and *Cck* (Veenvliet et al., 2013) and in the adult midbrain, expression can be detected in both the VTA and the SNc (**Figure 6D**). Similar to *Th,* no considerable differences could be observed between *Pitx3Cre/+; Ezh2* +/+ and *Pitx3Cre/+; Ezh2* L/L animals at both 3 months and 6 months of age. Together these results suggest that post-mitotic deletion of *Ezh2* does not influence the initial developmental programming of mdDA neurons, but that over time the expression of subset specific marks *Cck* and *Ahd2* are altered in a corresponding manner as observed in *En1Cre* driven *Ezh2* cKOs

### MdDA neurons are lost in aging animals lacking *Ezh2*

Previous studies have already shown that post-mitotic PRC2 deficiency can lead to delayed changes in expression of genes, including several important regulators of cell death (von Schimmelmann et al., 2016; Sun et al., 2014) and defective PRC2 activity has been associated with several degenerative diseases, including Huntington’s Disease (Li et al., 2013; von Schimmelmann et al., 2016; Seong et al., 2010; Södersten et al., 2014). We thus hypothesized that *Pitx3Cre* driven deletion of *Ezh2* might affects the survival of mdDA neurons next to programming defects as observed above. To establish whether cells are lost in *Pitx3/Ezh2* mutants we assessed the amount of TH+ neurons in the SNc and the VTA at two different ages, 3 months and 6 months, by means of immunohistochemistry (**Figure 7**). Quantification of the amount of TH+ cells in the VTA and the SNc of 3 month old *Pitx3Cre/+; Ezh2* +/+ (n=3) and *Pitx3/+; Ezh2* L/L (n=3) animals showed no significant differences in the amount of cells (**Figure 7A, B**). Interestingly, assessment of the amount of TH+ cells at 6 months of age showed a significant decrease in the amount of TH+ neurons in *Pitx3Cre/Ezh2* mutants (∼ 33% loss, n=3, p<0.05, two-tailed)(**Figure 7C**). Separate quantification of the cells in the VTA and SNc (**Figure 7A**, right panel, white dashed lines) demonstrated that only in the VTA of *Pitx3Cre/+; Ezh2* L/L animals a significant amount of TH+ cells is lost (∼32% loss, n=3, P<0.05, two-tailed) (**Figure 7A**, right panel, arrowhead, C), although a downward trend in the amount of TH+ in the SNc domain was detected (∼ 15% loss, n=3, p=0.07, two-tailed) (**Figure 7C**). The loss of cells in the VTA was confirmed by a loss of expression of *Calbindin D28K (Calb1),* a mark expressed by all cells of the VTA (**Supplemental figure 2,** arrowhead). Taken together our results demonstrate that post-mitotic deletion of *Ezh2* in mdDA neurons leads to a progressive loss of TH+ cells in the VTA, thereby supporting the argument that lower EZH2/ PRC2 activity may lead to neurodegeneration in specific neuronal populations.

**Figure 7:**
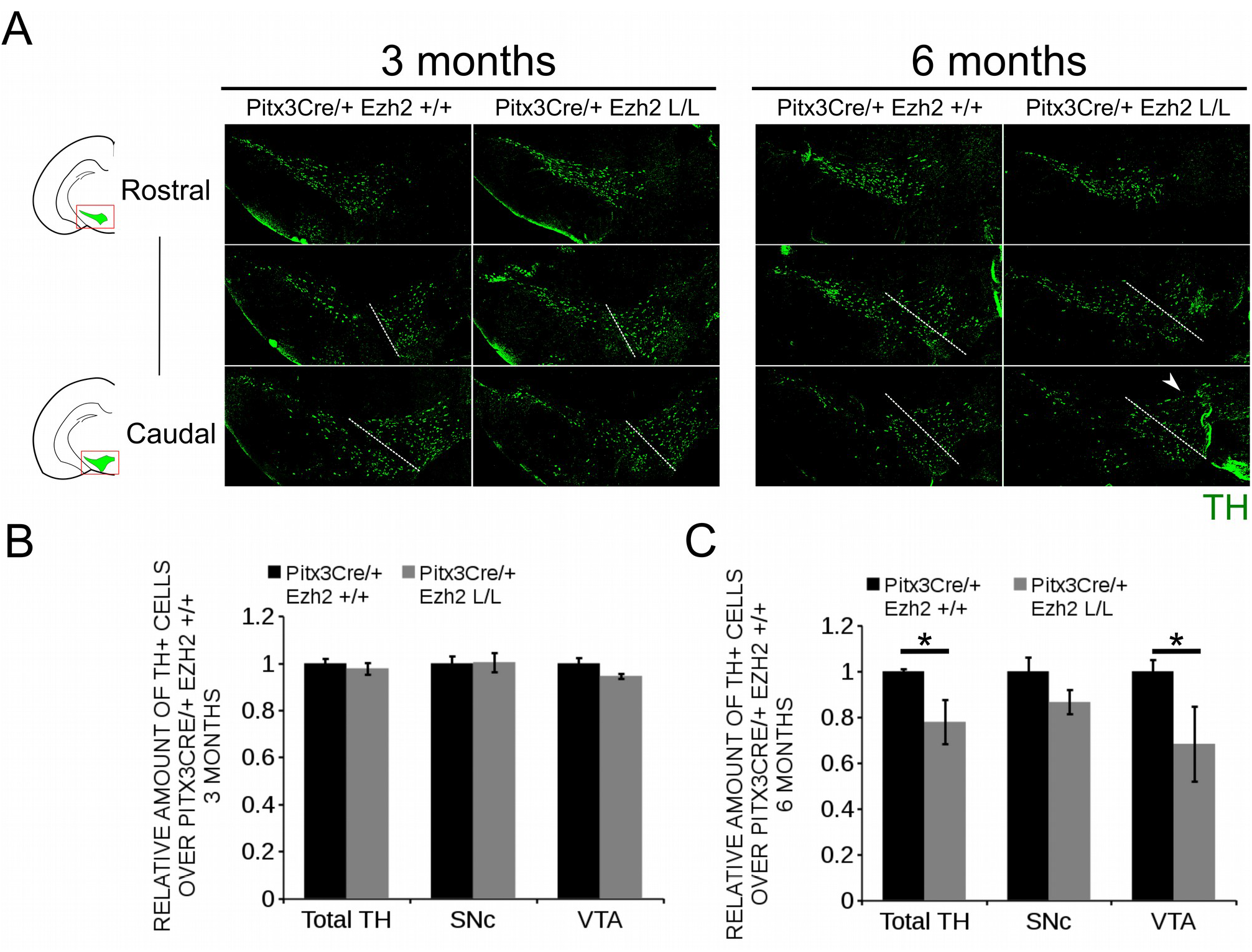
Progressive loss of TH+ cells in the VTA of 6 month old *Pitx3/+: Ezh2* L/L animals. (A) Immunohistochemistry for TH in coronal sections of the midbrain of 3 and 6 month old *Pitx3Cre/+; Ezh2 +/+* and *Pitx3Cre/+; Ezh2* L/L animals. Loss of TH+ neurons was observed in the VTA of 6 month old *Ezh2* cKO animals (white arrowhead). White dotted lines represent the border between what was considered SNc and VTA. (B) Quantification of the total amount of TH+ cells and the separation of the TH+ population into SNc and VTA TH+ neurons demonstrated no significant changes in either TH+ populations between 3 month old *Pitx3Cre/+; Ezh2* +/+ (black bars) and *Pitx3Cre/+; Ezh2* L/L (grey bars) animals (n=3). Wildtypes were set at 1. (C) Quantification of the total number of TH+ cells at 6 months of age showed a ∼22% reduction in the amount of TH+ cells in the *Ezh2* cKO (n=3, *P<0.05, two-tailed). Separations into the SNc and the VTA showed that most cells are lost in the VTA (∼ 32% loss, n=3, *P<0.05, two-tailed) and downward trend is observed in the amount of TH+ cells in the SNc (n=3, P=0.07, two-tailed). Wildtypes were set at 1.

### H3K27me3 is still present in mdDA neurons of 6 month old *Pitx3Cre* driven *Ezh2* cKOs

As described above, EZH2 functions as the methyltransferase of the PRC2 complex, which catalyzes the tri-methylation of H3K27 (Cao and Zhang, 2004b; Cao et al., 2002). The phenotype observed in *Pitx3Cre/+; Ezh2* L/L animals suggest that PRC2 activity might be altered, which could affect the presence of H3K27me3 in mdDA neurons of these animals. To verify whether the phenotype of *Pitx3Cre/+; Ezh2* L/L animals is a consequence of an overall loss of H3K27me3, we performed double immunohistochemistry for H3K27me3 and TH at 6 months of age (**Figure 8**). We analyzed the presence of H3K27me3 in TH+ neurons in both the SNc and the VTA. In the SNc H3K27me3 was present in TH+ neurons in *Pitx3Cre/+; Ezh2* L/L animals in a comparable manner to wildtype littermates (**Figure 8A (1,2,3,4)**). Similar results were obtained for the VTA, where H3K27me3 was also detected in TH+ cells of *Pitx3Cre/Ezh2* cKOs (**Figure 8B (1,2,3,4)**). Together our results show that post-mitotic deletion of *Ezh2* alone is not sufficient to remove overall H3K27me3 marks.

**Figure 8:**
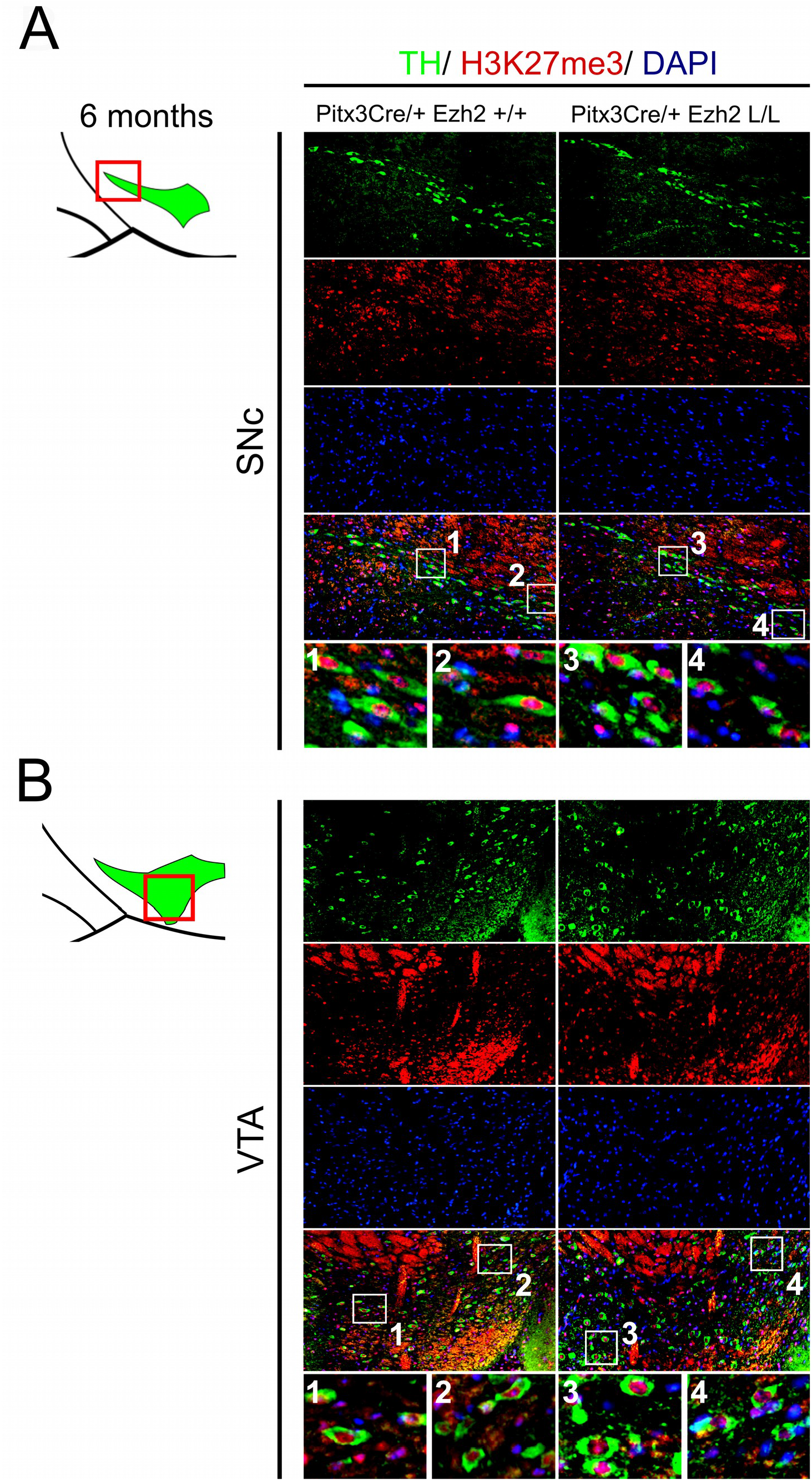
H3K27me3 can still be detected in the SNc and VTA of *Ezh2* cKO animals at 6 months of age. (A, B)Immunohistochemistry for TH (green) and H3K27me3 (red) in coronal sections of 6 month old midbrains demonstrated that H3K27me3 is still present in the SNc (A, 3,4))and VTA (B3,4)of *Pitx3Cre/+; Ezh2* L/L animals. Co-localization of H3K27me3 and DAPI staining was used to determine nuclear localization of the H3K27me3 staining.

### *Pitx3Cre/Ezh2* mutants display behavioral changes at 6 months of age

The different populations of mdDA neurons have characteristic projection areas and are involved in the regulation of different types of behavior. (Haber et al., 2000; Lynd-Balta and Haber, 1994; Prakash and Wurst, 2006). In *Pitx3Cre/+; Ezh2* L/L animals a loss of mdDA neurons is observed which might affect the dopaminergic output and alter the behavior of these animals. For this reason we assessed spontaneous locomotor activity in an open field test and climbing behavior of *Pitx3Cre/+; Ezh2* +/+ and *Pitx3Cre/+; Ezh2* L/L animals (**Figure 9**). An age-dependent effect on the climbing behavior was observed (**Figure 9A, B**). 6 month old *Ezh2* mutant mice (n=5) demonstrated a significantly lower climbing score than wildtype control animals (n=5) (∼ 20% lower, P<0.05, two-tailed) (**Figure 9A**), while in 3 month old *Pitx3Cre/; Ezh2* L/L animals climbing behavior was not affected (n=8) (**Figure 9A**). In contrast to the climbing behavior, spontaneous locomotor activity, assessed by distance walked and walking speed in an open field test, was not affected in both 3 month (n=8) and 6 month (n=5) old *Pitx3Cre/+; Ezh2* L/L animals in comparison to wildtype littermates (**Figure 9C, D**). These data demonstrate that the loss of *Ezh2* does not affect general locomotor activity, but specifically influences the more complex climbing behavior representing the loss of DA neurons in 6 months old *Pitx3Cre/+; Ezh2* L/L animals.

**Figure 9:**
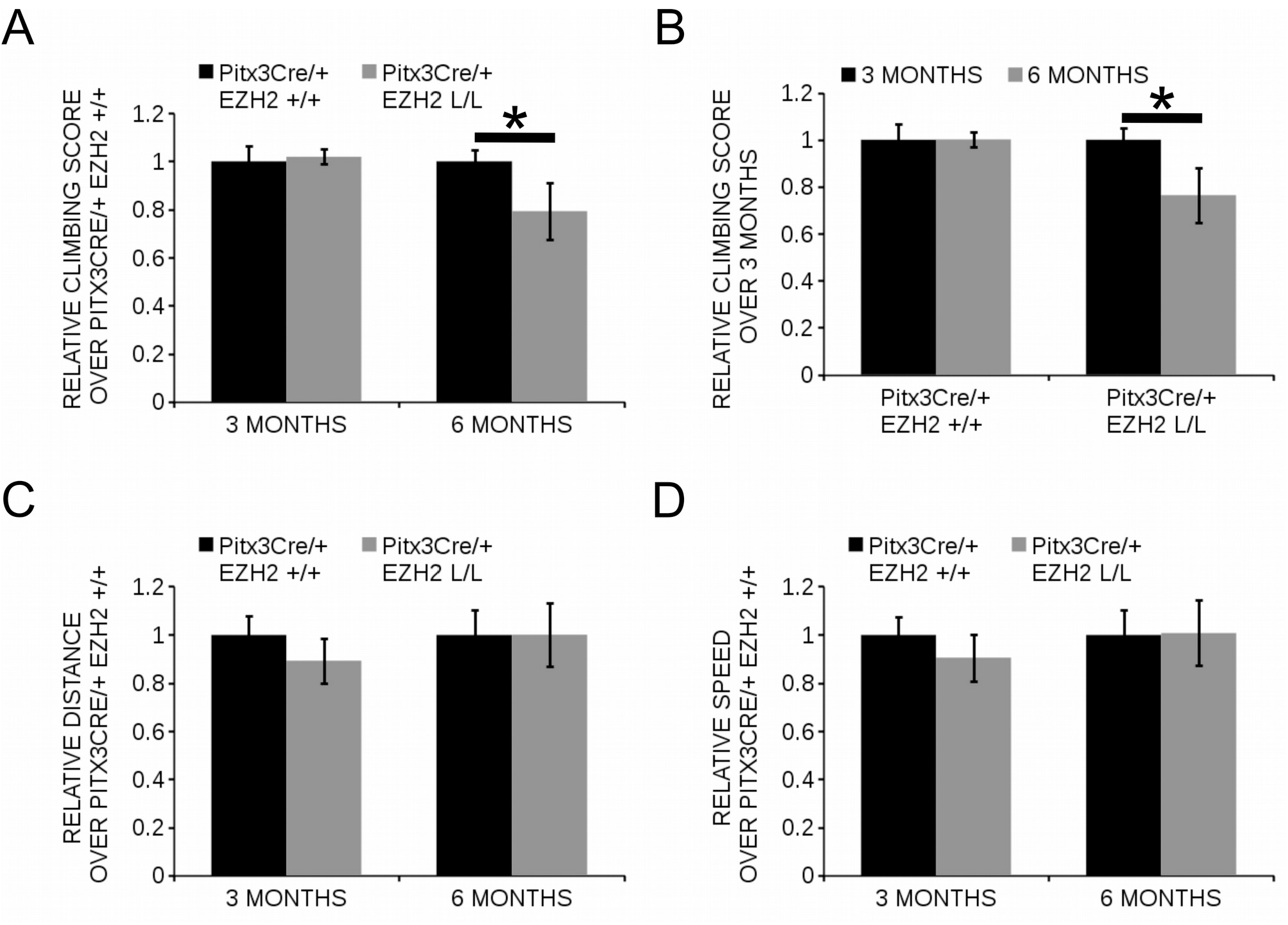
*Pitx3Cre* driven deletion of *Ezh2* influences climbing behavior of 6 month old animals. (A, B) Evaluation of climbing behavior at 3 months and 6 months of age. (A) 6 month old *Pitx3Cre/+;Ezh2* L/L (grey bars) animals have a significantly lower climbing score than wildtype littermates (black bars) (∼ 21% reduction, n=5, *P<0.05, two-tailed). This is not observed at 3 months of age (n=8). (B) Climbing-scores progressively reduce in the *Ezh2* cKO between 3 months (n=8, black bars) and 6 months (n=5, grey bars) (∼ 21% reduction, *P<0.05, two-tailed), while *Pitx3Cre/+;Ezh2* +/+ animals show similar climbing behavior at 3 months (n=8, black bars) and 6 months (n=5, grey bars). (C, D) Locomoter activity was monitored in an automated set up and the distance walked (in cm) was not significantly altered at 3 month (n=8) or 6 months (n=5) in *Pitx3Cre/+; Ezh2* L/L animals (C), nor was the speed (in cm/s) (D).

## Discussion

Over the last decade several protein have been identified to be differentially expressed in the SNc and the VTA (Chung et al., 2005). These differences in molecular profile have been implied to partially cause the selective degeneration of the SNc neurons in Parkinson’s disease, while neurons of the VTA remain unaffected (Oliveira et al., 2017). Individual subsets are dependent on different transcriptional programs for their development and maintenance (Hwang et al., 2003; Panman et al., 2014; Smidt et al., 2004; Smits et al., 2013; Veenvliet et al., 2013). In addition to the influence of transcription factors, recent studies have led to the hypothesis that modifications of histones might also influence the developmental program of neurons (Feng et al., 2016; Pereira et al., 2010; Zemke et al., 2015). H3K27me3 is a modification associated with gene-silencing and it shows a highly dynamic profile during development (Chou et al., 2011; Mikkelsen et al., 2007; Mohn et al., 2008; Yu et al., 2011). The methylation of H3K27me3 is catalyzed by the PRC2 complex, of which EZH2 is the methyltransferase (Cao and Zhang, 2004a). Previous studies in which *Ezh2* was conditionally ablated showed that *Ezh2* is involved during several stages of neurodevelopment (Di Meglio et al., 2013; Ezhkova et al., 2009; Feng et al., 2016; Pereira et al., 2010; Zemke et al., 2015). In the current study we focused on the role of *Ezh2* during the development, programming and maintenance of mdDA neurons. The conditional removal of *Ezh2* in early mdDA progenitors resulted in reduced amounts of TH-expressing cells at E14.5. Previous studies showed that neurogenesis is initiated earlier at the expense of self-renewal in embryos where *Ezh2* was conditionally removed from cortical neuroprogenitors (Pereira et al., 2010). Neurogenesis of mdDA neurons is initiated around E10.5 and ceases around E13.5, with the first TH+ neurons being detected around E11.5 (Bayer et al., 1995; Mesman et al., 2014). A shift in the balance between selfrenewal and differentiation, at the expanse of self-renewal, might explain why less TH+ neurons are present at E14.5 in the *Ezh2* cKO, however this is not reflected by the amount of neurons present at E12.5, as the number of TH+ cells are not increased at this time point. Another possible reason for the reduced number of TH+ neurons might be that mdDA progenitors and mdDA neurons are produced less efficient due to reduced canonical WNT signaling (Tang et al., 2009). In *Wnt1Cre/+; Ezh2* L/L embryos inhibitors of the WNT signaling pathway were up-regulated and less β-galactosidase-positive neural cells were detected in the dorsal midbrain (Zemke et al., 2015). In addition, analysis of *Wnt1* expression in *En1Cre/+; Ezh2* L/L embryos revealed reduced levels of *Wnt1* at E12.5 (Wever *et al.*, unpublished data), suggesting that WNT signaling is also affected in our model.

Interestingly, the loss of TH+ cells mostly affected the *Ahd2+* population, which is generated first according to birth-dating studies (Bayer et al., 1995; Bye et al., 2012), while the later born caudomedial population of *Cck+* cells remains largely unaffected. These results are in contrast with the results obtained by Pereira *et al.* where the earlier onset of cortical neurogenesis led to the loss of later born layer 2-4, due to a depletion of the neuroprogenitor pool (Pereira et al., 2010). The loss of the *Ahd2* population is also observed in *Pitx3* null mutants (Jacobs et al., 2007), however, in *En1Cre/+; Ezh2* L/L animals *Pitx3* levels are not affected and PITX3 expression mimicked TH, suggesting that the loss of the *Ahd2* positive population is not due to an effect of *Ezh2* on *Pitx3* expression. The impaired development of the rostrolateral population might be partially due to defective tangential migration. MdDA neurons are generated at the ventricular zone of the ventral midbrain and migrate to their final position first via radial migration followed by tangential migration (Kawano et al., 1995). The dislocated TH+ cells in the medial sections and the loss of rostrolateral cells, suggest that tangential migration might be affected in *En1Cre/+; Ezh2 L/L* embryos. Lateral migration is dependent on the interaction of mdDA neurons with tangential fibers (Kang et al., 2010). Reelin, an extracellular matrix glycoprotein, has been found to be essential for formation of these fibers, as genetic ablation of *Reelin* led to the loss of tangential fibers, while radial glial fibers were formed normally (Kang et al., 2010). A previous study demonstrated that genetic ablation of *Ezh2* in cortical progenitors affected neuronal migration by influencing *Reelin* expression (Zhao et al., 2015). In addition EZH2 was found to be required to maintain the tangential migratory program of pontine neurons (Di Meglio et al., 2013), hinting towards a role for EZH2 in tangential migration.

Besides a role for *Ezh2* in early development, we also demonstrated that *Ezh2* is important for the preservation of neuronal identity and the survival of a subset of TH+ neurons. Post-mitotic deletion of *Ezh2* leads to a progressive up-regulation of *Cck* in the rostral population of the mdDA system and a loss of *Ahd2* expression in the more caudal population. *Cck* has been found to be a target of PRC2 during development and loss of *Ezh2* in neuronal progenitors leads to a significant up-regulation of *Cck* expression (Bracken et al., 2006; Ku et al., 2008; Mikkelsen et al., 2007; Pereira et al., 2010). We thus theorize that the progressive appearance of *Cck* in the rostral population of 6 month old *Pitx3/Ezh2* mutants is because H3K27me3 on the *Cck* promoter is not maintained over time, leading to the de-repression of the promoter. In contrast, the promoter of *Ahd2* has not been associated with EZH2 binding and H3K27me3 (Mikkelsen et al., 2007), suggesting that the loss of *Ahd2* in the caudal mdDA system is not due to a direct effect of EZH2 on *Ahd2* expression. In addition to alterations in expression of subset marks, *Pitx3Cre/Ezh2* mutants also display a progressive loss of TH+ cells in the VTA. The loss of TH+ cells was reflected by reduced expression of the VTA subset mark, *Calb1*. In contrast to *Cck, Calb1* is expressed by all neurons of the VTA (Brignani and Pasterkamp, 2017; La Manno et al., 2016), suggesting that *Pitx3Cre* driven deletion of *Ezh2* specifically affects the *Calb+Cck-* population of VTA neurons. Calb1 has been shown to promote resistance against neurodegeneration (German et al., 1992; McMahon et al., 1998) and the reduced levels of *Calb1* in the more caudal region of the mdDA system might contribute the loss of TH+ neurons, however the loss of *Calb1* might also be a consequence of the initial loss of neurons. Next to a loss in DA neurons, *Pitx3Cre/+; Ezh2* L/L also showed reduced climbing behavior without affecting general locomotor activity. It is hypothesized that climbing behavior requires different and more complex dopaminergic mechanisms than horizontal locomotor behavior (Cabib and Puglisi-Allegra, 1985; Moore and Axton, 1988; Usiello et al., 2000), including projections from the VTA to the Nucleus accumbens (Costall et al., 1983, 1985; Salamone, 1992), which might explain the specific reduction in climbing behavior. However, it needs to be noted that *Pitx3* is also expressed in the muscles and lens and *Pitx3Cre* driven deletion of *Ezh2* may also affect these systems. In addition, *Pitx3Cre/+; Ezh2* L/L animals had a higher change of dying prematurely (∼35% higher).

Even though both *En1/Ezh2* and *Pitx3/Ezh2* mutants display a disturbance in *Cck* and *Ahd2* expression and a loss of TH+ cells, the severity of the phenotype differs. The phenotype observed in *En1Cre/+; Ezh2* L/L embryos is probably a consequence of an overall defect in PRC2 functioning and a global loss of H3K27me3 in the midbrain area. In contrast, the effects observed in *Pitx3Cre* driven *Ezh2* cKOs can not be explained by a general lack of H3K27Me3, as the mark was still detected in the SNc and VTA of 6 month old *Pitx3Cre/+; Ezh2* L/L animals. However, EZH2 has also been shown to have PRC2-independent activity and active transcription of genes marked by H3K27me3 has also been observed before, suggesting that changes in *Ezh2* functioning might not be reflected by changes in H3K27me3 (Bracken et al., 2006; Mikkelsen et al., 2007; Mohn et al., 2008; Xu et al., 2012). In addition, local changes in H3K27me3 levels are not visualized by immunohistochemistry for H3K27me3.

Taken together, this study shows that next to a role in development, *Ezh2* is also important for the maintenance and survival of cells of a small group of mdDA neurons. Interestingly, the loss of *Ezh2* mainly affects the embryonic development of the rostrolateral population, destined to become the SNc, while cells of the VTA are lost when *Ezh2* is removed post-mitotically. This suggests that *Ezh2* has different functions during development and neuronal maintenance.

## Acknowledgements

We would like to thank Dr. Frederic Zilbermann of the Friedrich Miescher Institute for his generous gift of the *Ezh2*-floxed mouse line and Dr. Sandra Blaess for kindly providing us with the *En1Cre; R26RYFP* mouse strain. This work was sponsored by the NWO-ALW (Nederlandse Organisatie voor Wetenschappelijk Onderzoek-Aard en Levenswetenschappen) VICI grant (865.09.002) awarded to Prof Dr. Marten P. Smidt.

---

**Supplemental Figure 1:**
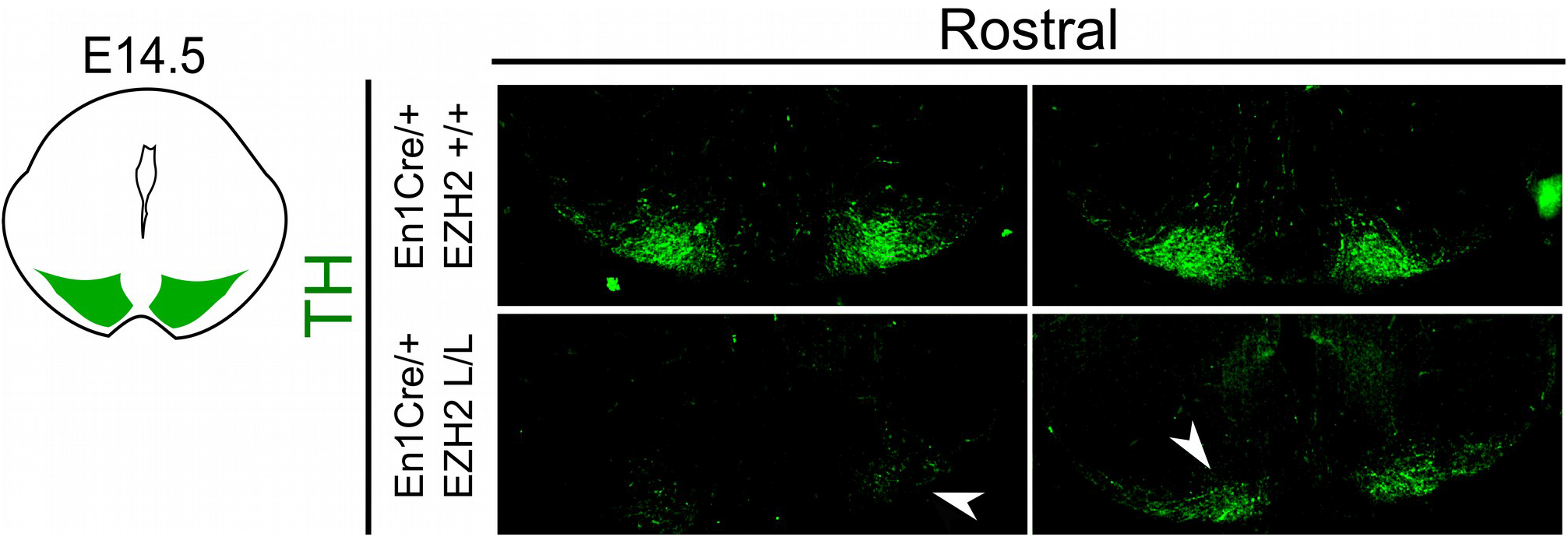
TH expression is reduced in rostral sections of the embryonic midbrain of *En1/Ezh2* mutants. Analysis of TH expression in coronal E14.5 midbrain sections by means of immunohistochemistry. Expression of TH is reduced in the rostral sections (white arrowheads).

**Supplemental Figure 2:**
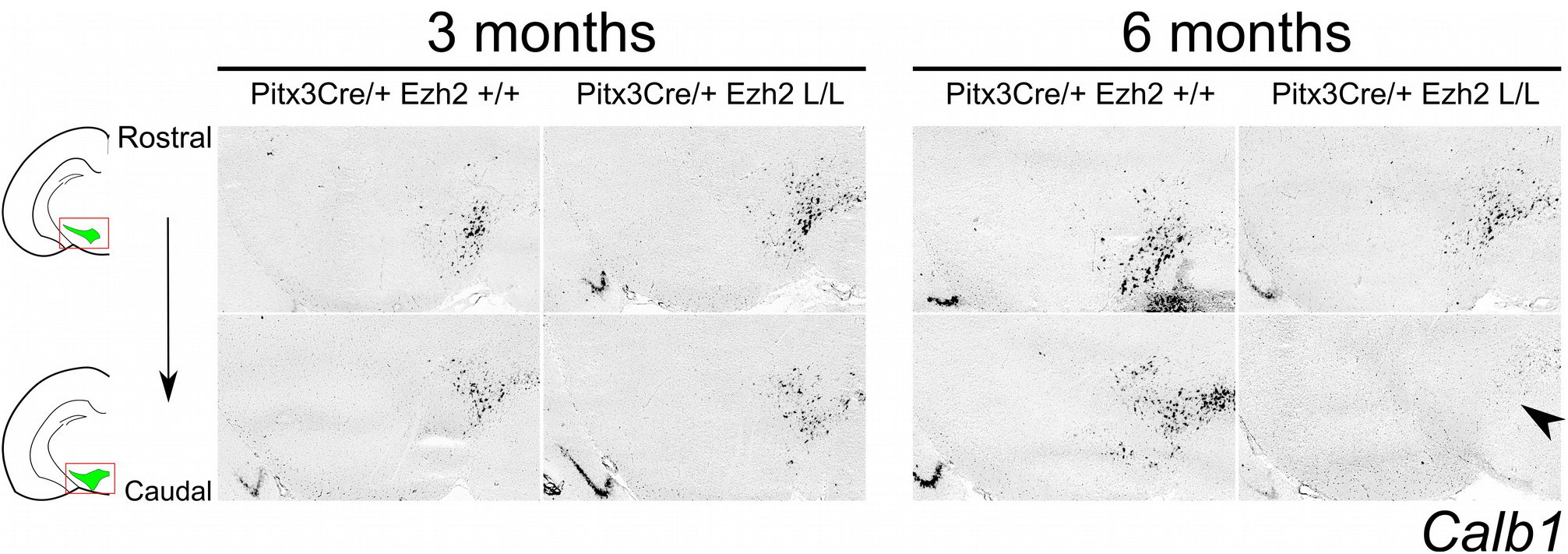
*Pitx3Cre* driven deletion of *Ezh2* induces a progressive loss of *Calb1* staining in the VTA. Analysis of the expression of *Calb1* in coronal midbrain sections of 3 month and 6 month old *Pitx3Cre/+; Ezh2* +/+ and *Pitx3Cre/+; Ezh2* L/L animals by means of *in situ* hybridization. Expression of *Calb1* is lost in the caudal VTA of 6 month old *Ezh2* cKO animals (black arrowhead), while expression can still be detected in this location at 3 months (left panel).

